# Spatiotemporal chloride dynamics in hypothalamic CRH^PVN^ neurons

**DOI:** 10.1101/2020.05.29.122432

**Authors:** Aaron Lanz, Grant Gordon, Jaideep Bains

## Abstract

Chloride (Cl^-^) dynamics shape inhibitory neurotransmission in the brain. In the hypothalamus, GABA signalling onto corticotropin-releasing hormone (CRH^PVN^) neurons can switch from inhibitory to excitatory during stress. Although Cl^-^ fluctuations mediate this stress-dependent shift in GABA signalling, the underlying Cl^-^ dynamics are poorly understood. Here, using a novel optogenetic strategy to load intracellular Cl^-^ using halorhodopsin, we demonstrate that KCC2 rapidly restores Cl^-^ setpoints in CRH^PVN^ neurons from naïve animals, but that this process is slowed following stress. Further, we report that somatic Cl^-^ homeostasis remains intact after stress. Our results strongly support the idea that KCC2 functions primarily to maintain Cl^-^ setpoints and that inhibitory synapses onto dendritic and somatic compartments of CRH^PVN^ neurons are differentially regulated during stress.

## Introduction

The functional organization of inhibition throughout the brain shapes how circuits process and store information. Inhibitory synaptic signalling can vary considerably across space, depending on the presynaptic cell type^1^, postsynaptic cell-compartment^1–3^, and local anionic environment^4,5^; and across time, depending on age^6^, experience^7^, and behavioural state^8,9^. Owing to the tight relationship between Cl^-^ and inhibition, much of this heterogeneity arises from variations in the factors that establish and maintain Cl^-^ gradients. Failures to regulate Cl^-^ levels contribute to several nervous system pathologies including chronic pain^10–12^, chronic stress^13^, schizophrenia^14^, autism^15^, and epilepsy^16–18^.

Corticotropin-releasing hormone neurons of the paraventricular nucleus of the hypothalamus (CRH^PVN^) gate the hormonal^19^ and behavioural^19–21^ responses to stress. In the absence of stress, the activity of CRH^PVN^ neurons is curtailed by robust inhibitory tone. In response to stress, there is a functional downregulation of the cation-Cl^-^ cotransporter KCC2 in CRH^PVN^ neurons to promote Cl^-^ accumulation during repetitive synaptic stimulation^9^. This shift in Cl^-^ extrusion capacity contributes to excitatory GABA signalling and the liberation of stress hormones^9,22^. Although we have identified a link between KCC2, Cl^-^, and inhibition during stress, we know little about the underlying Cl^-^ dynamics. Experimental and theoretical work suggests that KCC2 can buffer Cl^-^ loads^8,9,23^ and restore Cl^-^ gradients following equilibrium setpoint deviations^2,8,9,23–26^. Further, KCC2 expression and Cl^-^ regulation often vary between neuronal compartments^2,27,28^. The relative contribution of KCC2 to Cl^-^ buffering and restoration, and how these processes interact with synaptic location and stress is not known. Answering these questions is critical for understanding how inhibition shapes CRH^PVN^ physiology in the absence and presence of stress. For example, if KCC2 heavily contributes to Cl^-^ setpoint restoration, then KCC2 downregulation during stress may promote substantial fluctuations in the magnitude and direction of GABA signalling^29^. Furthermore, if Cl^-^ is differentially regulated in dendritic and somatic compartments, then GABA inputs to different locations on the cell will have distinct effects on CRH^PVN^ physiology.

A major barrier limiting the study of KCC2 and Cl^-^ dynamics is the absence of simple tools to directly measure Cl^-^ homeostasis. Many studies have relied on gramicidin patch clamp recordings of GABA currents, assuming that a depolarization in the Cl^-^ equilibrium potential reflects a downregulation in KCC2^6,8,9,12,13,17,18,30,31^. However, this assumption is not necessarily valid; blockade of KCC2 can elevate or reduce Cl^-^ levels depending on the initial Cl^-^ concentration^4,5^, which itself can vary considerably across and within cell populations^4,5^. Furthermore, these measurements are relatively slow and cannot reveal fast Cl^-^ dynamics. To circumvent these issues, some groups have measured Cl^-^ dynamics by tracking GABA currents during high intensity GABA_A_ stimulation^2,8,9,23,26,32^. Although these approaches revealed aspects of Cl^-^ homeostasis, the use of GABA receptors to measure and manipulate Cl^-^ levels can complicate data interpretation.

Here, we used an optogenetic tool along with whole-cell recordings from identified CRH^PVN^ neurons in hypothalamic brain slices to discern chloride dynamics and better understand the consequences of acute stress. We report that KCC2 rapidly restores Cl^-^ setpoints in CRH^PVN^ neurons but does not influence the Cl^-^ buffering capacity of inhibitory synapses. Cl^-^ buffering is instead tightly linked to synaptic location. Furthermore, we demonstrate that stress slows Cl^-^ setpoint restoration at dendritic synapses, but not somatic compartments, suggesting a differential role of dendritic and somatic synapses in CRH^PVN^ physiology during stress.

## Methods

### Animals

All animal protocols were approved by the University of Calgary Animal Care and Use Committee (AC14-0138). Experiments were performed on 6- to 8-week-old male and female *Crh-IRES-Cre; Ai14* mice, in which CRH neurons express Cre and tdTomato fluorophore. Mice were individually housed on a 12 h/12 h light/dark cycle (lights on at 7:00 hours) and were given ad libitum access to food and water.

### AAV Virus Injection

6- to 8-week-old *Crh-IRES-Cre; Ai14* mice were placed into a stereotaxic apparatus under isoflurane anaesthesia. Pulled glass capillaries, filled with recombinant adeno-associated virus (AAV) carrying enhanced halorhodopsin 3.0 (AAV2/9-EF1α-DIO-eNpHR3.0-eYFP, Institut Universitaire en santé mentale de Montréal, lot 1155), were inserted into the brain at coordinates slightly dorsal to the left PVN (anteroposterior (AP), 0 mm; lateral (L), 0.2 mm from the bregma; dorsoventral (DV), −4.7 mm from the dura). Virus was pressure injected into the brain three times with a Nanoject II apparatus (Drummond Scientific Company), each injection at a volume of 69 nL and placed increasingly deeper into the brain (AP: −4.7 mm, −4.8 mm, −4.9 mm). Injections were repeated for the right PVN (AP, 0 mm; L, −0.2 mm from the bregma; DV, −4.7 mm from the dura). Following surgery, animals were allowed to recover for at least two weeks prior to electrophysiology experiments.

### Slice Preparation

Naïve or acutely stressed mice (30-minute restraint inside a 50 mL falcon tube) were anesthetized with isoflurane and decapitated. Brains were rapidly dissected and immersed in cold artificial cerebrospinal fluid (aCSF) (0-4 °C, 95% O_2_/5% CO_2_ saturated) containing (in mM) 126 NaCl, 2.5 KCl, 26 NaHCO_3_, 2.5 CaCl_2_, 1.5 MgCl_2_, 1.25 NaH_2_PO_4_, and 10 glucose. Coronal slices (250 μm), obtained using a Leica vibratome, were incubated in an N-Methyl-D-glucamine (NMDG) recovery solution (37 °C, 95% O_2_/5% CO_2_ saturated) containing 110 NMDG, 2.5 KCl, 25 NaHCO_3_, 0.5 CaCl_2_, 10 MgCl_2_, 1.2 NaH_2_PO_4_, 25 glucose, and 110 HCl (pH = 7.5) for 15 minutes and then warm aCSF (37 °C, 95% O_2_/5% CO_2_ saturated) for 45+ minutes. Slices were removed from heat and remained in room temperature aCSF until use.

### Corticosterone Immunoassay

During slice preparation, trunk blood was collected into cold EDTA-coated tubes and then centrifuged (8000 r.p.m., 4 °C, 20 min) to separate the plasma and blood cells. Aliquots of plasma were stored at −20 °C and then assayed with DetectX Corticosterone Immunoassay kits (Arbor Assay) at a later date. Each plasma sample was assayed three times to obtain an average value for each animal.

### Electrophysiological Recordings

Slices were transferred to a recording chamber superfused with aCSF (1 mL/min, 30-32 °C, saturated with 95% O_2_/5% CO_2_) and neurons were visualized using an upright microscope (BX51WI, Olympus) fitted with differential interference contrast optics, epifluorescence optics (BH2-RFL-T3, Olympus), and a digital camera (IR-1000 DAGE-MTI). Borosilicate glass pipettes (3-5 MΩ) were filled with a KGluconate internal solution containing (in mM): 108 gluconate, 8 KCl, 8 Na-gluconate, 1 K_2_EGTA, 10 HEPES, 2 MgCl_2_, 4 K_2_ATP, and 0.3 Na_3_GTP for whole cell recordings. Recordings were made of CRH^PVN^ neurons by selectively targeting tdTomato-expressing cells adjacent to the III ventricle. Halorhodopsin-mediated photocurrents were elicited by voltage-clamping cells at −30 mV and illuminating the PVN with 595 nm yellow light through a fiber-coupled LED (M595F2, Thorlabs). Cells with photocurrents below 30 pA were typically discarded. GABA currents (I_GABA_) were elicited in one of two ways: pressure application of a GABA solution (10 μM in aCSF; Tocris #5A/205401) using a Picospritzer II (General Valve Corporation); or electrical stimulation of GABA synapses using a monopolar electrode (100-150 μs, 10-60 V, S44 stimulator, Grass instruments) placed in an aCSF-filled borosilicate glass pipette (3-5 MΩ) in the presence of 6,7-Dinitroquinoxaline-2,3-dione (DNQX; 10 μM; Tocris #14A/166549).

The reversal potential of evoked inhibitory postsynaptic currents (eIPSCs) was obtained using a brief (20 s) voltage step protocol. The voltage was held at −70 mV for 1.29 s, stepped towards a test potential for 1.42 s, during which an eIPSC was evoked, and then returned to −70 mV for 1.29 s. The protocol was repeated five times to obtain an eIPSC at −90, mV, −80 mV, −70 mV, −60 mV, and −50 mV test potentials.

During Cl^-^ homeostasis experiments, cells were voltage clamped at −30 mV and I_GABA_ was elicited at 1 Hz. I_GABA_ was triggered for 40 s without light illumination, 40 s with light illumination, and then 80 s without light illumination. In some experiments, VU 0240551 (3888, Tocris), dissolved in DMSO (100 mM), was added to the circulating aCSF to a final concentration of 10 μM.

Access resistance was monitored every three minutes, and recordings were discarded if the resistance exceeded 20 MΩ or changed by >15%.

### Data Analysis

Signals were amplified using a Multiclamp 700B amplifier (Molecular Devices), low-pass filtered at 1 kHz, digitized at 10 kHz using the Digidata 1440 (Molecular Devices), and recorded for offline analysis using pClamp 10.2 (Molecular Devices).

I_GABA_ amplitude was measured by comparing the peak change in current following GABA pressure application or electrical stimulation. The eIPSC reversal potential (E_IPSC_) was calculated by quantifying the voltage where eIPSC reaches 0 pA. For Cl^-^ homeostasis experiments, the first 20 I_GABA_ were discarded and then the second 20 I_GABA_ were used as a baseline for normalization. The collapse was analyzed in two ways: the rate of collapse and the magnitude of collapse. The rate of collapse was calculated by fitting a monoexponential curve to the I_GABA_ during light illumination and extracting the tau value:

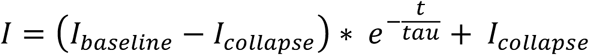

Where *I*_*baseline*_ is the I_GABA_ amplitude at baseline, *I*_*collapse*_ is the I_GABA_ amplitude at the end of the collapse, *t* is the time, and *tau* is the time when I_GABA_ amplitudes have reached 63.21% of the collapse value. The magnitude of collapse was calculated as the percent change in I_GABA_ amplitude during the last 20 seconds of light illumination in comparison to the baseline:

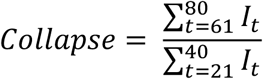

Where *I*_*t*_ is the I_GABA_ amplitude at time = t. The recovery of Cl^-^ currents was determined by calculating the rate of recovery. A monoexponential curve was fit to the I_GABA_ following light stimulation and extracting the resulting tau value:

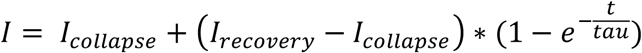

Where *I*_*collapse*_ is the I_GABA_ amplitude at the end of the collapse, *I*_*recovery*_ is the I_GABA_ amplitude at the end of recovery, *t* is the time, and *tau* is the time when I_GABA_ amplitude has reached 63.21% of the recovery value. These functions were generated and evaluated using GraphPad Prism 8. Finally, the photocurrent and eIPSC slope were obtained for additional analysis. During light illumination, the light would turn off briefly (31.2 ms) before each I_GABA_. The photocurrent could be measured for each cycle by comparing the current during this brief off-period to the current immediately following. The average photocurrent over the 40 s light illumination period was used for all photocurrent analyses. The eIPSC slope was obtained by measuring the maximum rate of change during the rising phase of synaptic currents.

### Statistics

The normality of all data was checked using the D’Agostino & Pearson omnibus normality test with an a = 0.05. When normality could not be assumed, nonparametric tests were used. For comparisons between two groups, a two-tailed nonparametric Mann-Whitney U test was used with a = 0.05. For comparisons between multiple groups, a two-tailed Friedman test was used with a = 0.05. Significant differences on the Friedman test were followed up by Dunn’s multiple comparison test. When normality could be assumed, parametric tests were used. For comparisons between two groups, a two-tailed unpaired t-test was used with a = 0.05. The results were the same when nonparametric or parametric tests were used. All statistical tests were run on GraphPad Prism 8. All data was presented as mean ± standard error of the mean. Curve fitting was done on SPSS using the curve estimation function. The relationship between two variables was fitted by a set of functions, including linear, inverse, exponential, quadratic, cubic, and logarithmic. The fit of each curve was evaluated using an *R*^*2*^ value, which indicates how much of the variance is explained by the curve, and by an ANOVA, which indicates the probability of obtaining the data if the two variables are not related by the curve. The ANOVA was run with a = 0.05.

## Results

### An optogenetic strategy to study Cl^-^ homeostasis

We measured Cl^-^ dynamics in CRH^PVN^ neurons across different neuronal compartments in the presence or absence of stress. Previous attempts have relied on electrophysiological measurements of GABA or inhibitory synaptic currents while manipulating Cl^-^ gradients by applying inhibitory neurotransmitters^2,25,32^ or repetitively stimulating inhibitory synapses^2,8,9,33^. These approaches are limited because the techniques used to measure and perturb Cl^-^ gradients are interdependent; that is, high intensity stimulation of GABA_A_ receptors or inhibitory synapses can interfere with inhibitory current measurements by eliciting receptor desensitization^2,34^ or short-term plasticity^35^. Here, we used the light-sensitive Cl^-^ pump, enhanced halorhodopsin, to elevate intracellular Cl^-^ levels with light^36^, and thus decouple Cl^-^ manipulation from Cl^-^ measurement. We expressed halorhodopsin in CRH^PVN^ neurons by injecting a cre-dependent *AAV-DIO-eNphR3*.*0-eYFP* virus into the PVN of *CRH-IRES-cre; Ai14* transgenic animals^37^ (**Figure 1a,b**). Two weeks after virus injection, illumination of CRH^PVN^ neurons in hypothalamic brain slices with 595 nm light resulted in outward currents (photocurrent magnitude: 30-150 pA) that robustly inhibited CRH^PVN^ neuron spiking (**Figure 1c**). We measured the reversal potential of evoked inhibitory synaptic currents (eIPSCs) before, during, and after halorhodopsin activation (**Figure 1d**). Within 20 seconds of light onset, the average eIPSC reversal potential depolarized and then recovered within 45 seconds of light termination (pre-light: −69.1 ± 2.0 *n* = 10, *N* = 9; light: −58.6 ± 1.1, *n* = 10, *N* = 9; post-light: −70.4 ± 2.1, *n* = 10; Friedman test, *Q* = 15.2, *p* < 0.0001; **Figure 1d-f**). This corresponds to a 4.5 mM average elevation in intracellular Cl^-^. Thus, halorhodopsin activation can transiently elevate intracellular Cl^-^ levels in CRH^PVN^ neurons.

**Figure 1.**
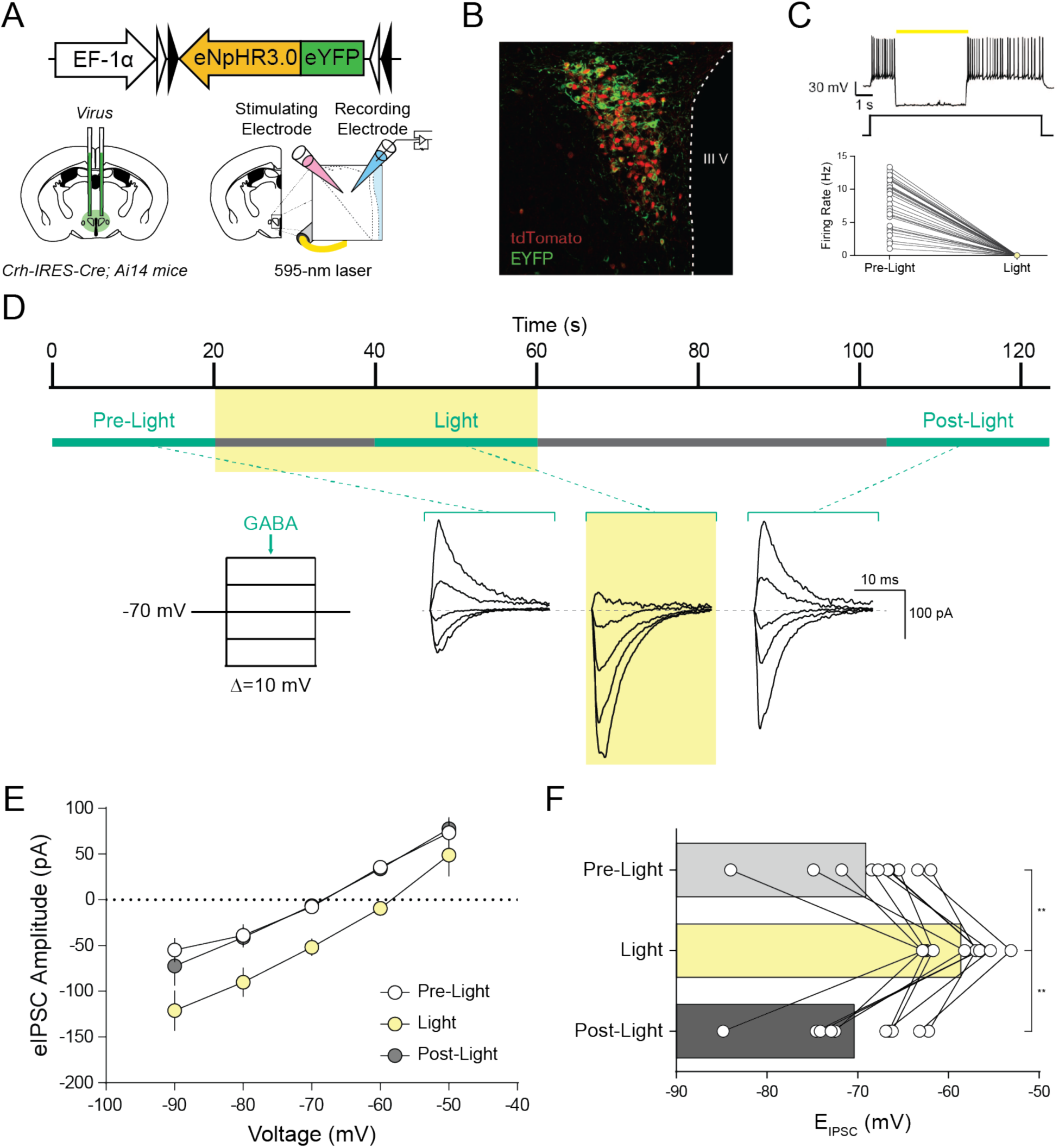
Halorhodopsin activation transiently shifts E_IPSC_. (A) Top, Cre-dependent *AAV-DIO-eNphR3*.*0-eYFP* virus injected into the PVN. Bottom left, halorhodopsin virus was injected bilaterally into the PVN of *CRH-IRES-Cre; Ai14* mice. Bottom right, schematic of electrode and laser configuration for electrophysiological study of CRH^PVN^ neurons (red). (B) Confocal image showing expression of eYFP (green, labelling eNphR3.0) and tdTomato (red, labelling CRH neurons) in hypothalamic brain slices. III V represents the third ventricle. (C) Top, Inset, action potentials during a 10-30 pA current step before, during (yellow bar), and after 595 nm light illumination. Bottom, halorhodpsin activation abolished current-mediated spiking in CRH^PVN^ neurons. (D) Top, timeline of experiment. E_IPSC_ was determined using a brief 20 s voltage step protocol (green lines) before, during, and after halorhodopsin activation. Yellow bar represents 595 nm light illumination. Bottom, representative example of inhibitory synaptic currents elicited at various holding potentials (−90 mV to −50 mV, at 10 mV increments) before, during, and after light illumination. (E) Current-voltage plot of eIPSCs before, during, and after halorhodopsin activation. (F) Halorhodopsin activation transiently depolarized the E_IPSC_ of inhibitory synapses on CRH^PVN^ neurons (Friedman test, *Q* = 15.2, *p* < 0.0001).

The restoration of Cl^-^ setpoints by homeostatic machinery can happen on the order of seconds^2,23,32^. To detect fast Cl^-^ dynamics during halorhodopsin activation, we voltage-clamped cells at −30 mV and stimulated inhibitory synapses at a frequency of 1 Hz. By clamping the voltage at −30 mV, we could use fluctuations in eIPSC amplitudes to track relative changes in Cl^-^ gradients (Figure 2a). Indeed, eIPSC amplitudes decayed to a new steady state during halorhodopsin-mediated Cl^-^ loading, and then rapidly recovered after light termination (**Figure 2a**). Both the collapse and recovery of Cl^-^ gradients during Cl^-^ homeostasis were fit well with mono-exponential functions, allowing us to approximate the time constant (tau) of Cl^-^ gradient decay and recovery. The average tau of eIPSC collapse during halorhodopsin activation was 4.3 ± 0.87 s (mono-exponential decay, *R*^*2*^ = 0.51 ± 0.04, **Figure 2b**), while the average tau of eIPSC recovery was 3.9 ± 0.89 s (mono-exponential association, *R*^*2*^ = 0.34 ± 0.04, **Figure 2c**). In addition, we could calculate the degree of Cl^-^ collapse by comparing the normalized amplitudes of eIPSCs at the end of light illumination with baseline values. During halorhodopsin activation, eIPSC amplitudes collapsed to 48.8 ± 6.9% of baseline values (**Figure 2b**). We did not explicitly measure the percent recovery of Cl^-^ gradients because synaptic currents recovered to near baseline values in almost all experiments. Thus, halorhodopsin and voltage-clamp electrophysiology can be used as a strategy to quantify synaptic Cl^-^ homeostasis in CRH^PVN^ neurons.

**Figure 2.**
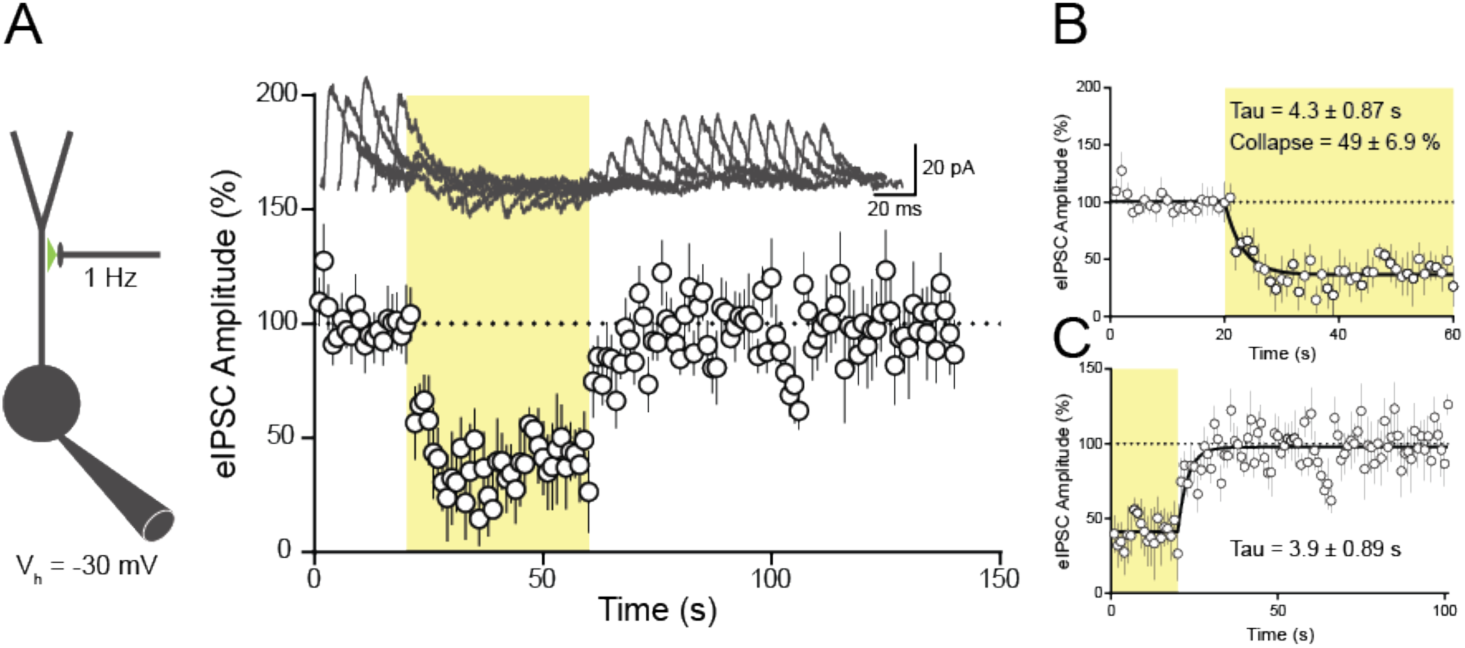
Temporal kinetics of synaptic Cl^-^ homeostasis in CRH^PVN^ neurons. **(A)** Left, schematic of stimulation and recording configuration. CRH^PVN^ neurons were voltage clamped at −30 mV and eIPSCs were elicited at 1 Hz. Right, amplitude of eIPSCs before, during, and after halorhodopsin activation, normalized to pre-light amplitudes. Inset, representative example of eIPSCs throughout experiment. **(B)** Average eIPSC amplitudes before and during light illumination, normalized to pre-light amplitudes. Data is fit with a monoexponential decay function (mean *R*^*2*^ = 0.51 ± 0.04) with an average tau of 4.3 ± 0.87 s and collapse of 48.8 ± 6.9. **(C)** Average eIPSC amplitudes during and after light illumination, normalized to pre-light amplitudes. Data are fit with a monoeponential association function (*R*^*2*^ = 0.34 ± 0.04) with an average tau of 3.9 ± 0.89 s. Yellow bars represent 595 nm light illumination.

### KCC2 does not set the Cl^-^ buffering capacity of CRH^PVN^ neurons

Endogenous homeostatic mechanisms, such as KCC2 cotransporters, may allow synapses to buffer Cl^-^ loads. In CRH^PVN^ neurons and VTA interneurons, KCC2 downregulation facilitates the accumulation of Cl^-^ during repetitive GABA synaptic stimulation^8,9^, while in hippocampal pyramidal neurons, KCC2 overexpression impedes Cl^-^ rundown^23^. To measure the influence of KCC2 on the Cl^-^ buffering capacity of inhibitory synapses on CRH^PVN^ neurons, we measured the tau and magnitude of halorhodopsin-mediated Cl^-^ accumulation in the presence of the specific KCC2 inhibitor, VU 020551^38–40^. Vehicle-treated (0.01% DMSO) cells and untreated cells had identical Cl^-^ dynamics and were thus pooled into one control group for all analyses. Pharmacological blockade of KCC2 using VU 020551 failed to alter the rate of eIPSC collapse during halorhodopsin activation (control mean tau = 4.3 ± 0.87 s, *n* = 17, *N* = 10; VU 020551 mean tau = 5.1 ± 1.2 s, *n* = 8, *N* = 5; Mann-Whitney test, *U* = 49.0, *p* = 0.29; **Figure 3a**), or the magnitude of eIPSC collapse (control mean = 48.8 ± 6.9 %, *n* = 18, *N* = 11; VU 020551 mean= 21.9 ± 15.7 %, *n* = 9, *N* = 6; Mann-Whitney test, *U* = 47.0, *p* = 0.08; **Figure 3b**). The failure of VU 020551 to impact the collapse of Cl^-^ gradients was not due to differences in photocurrents (control mean = 67.5 ± 6.7 pA, *n* = 18, N = 11; VU 020551 mean = 61.3 ± 12.0 pA, *n* = 9, *N* = 6; Mann-Whitney test, *U* = 66.0, *p* = 0.45; **Figure 3c**) or synapse location as approximated using the eIPSC slope^41^ (control mean = 70.2 ± 11.8 pA/ms, *n* = 17, *N* = 10; VU 020551 mean = 80.6 ± 23.6 pA/ms, *n* = 8, *N* = 5; Mann-Whitney test, *U* = 65.0, *p* = 0.88; **Figure 3d**). A simple explanation for this observation is that halorhodopsin photocurrents overwhelmed KCC2-mediated Cl^-^ extrusion, even when photocurrents were relatively small, and as a result, the degree of Cl^-^ collapse was constrained by other factors, such as cell compartment volume^2^ or synapse proximity to the patch pipette^42^.

**Figure 3.**
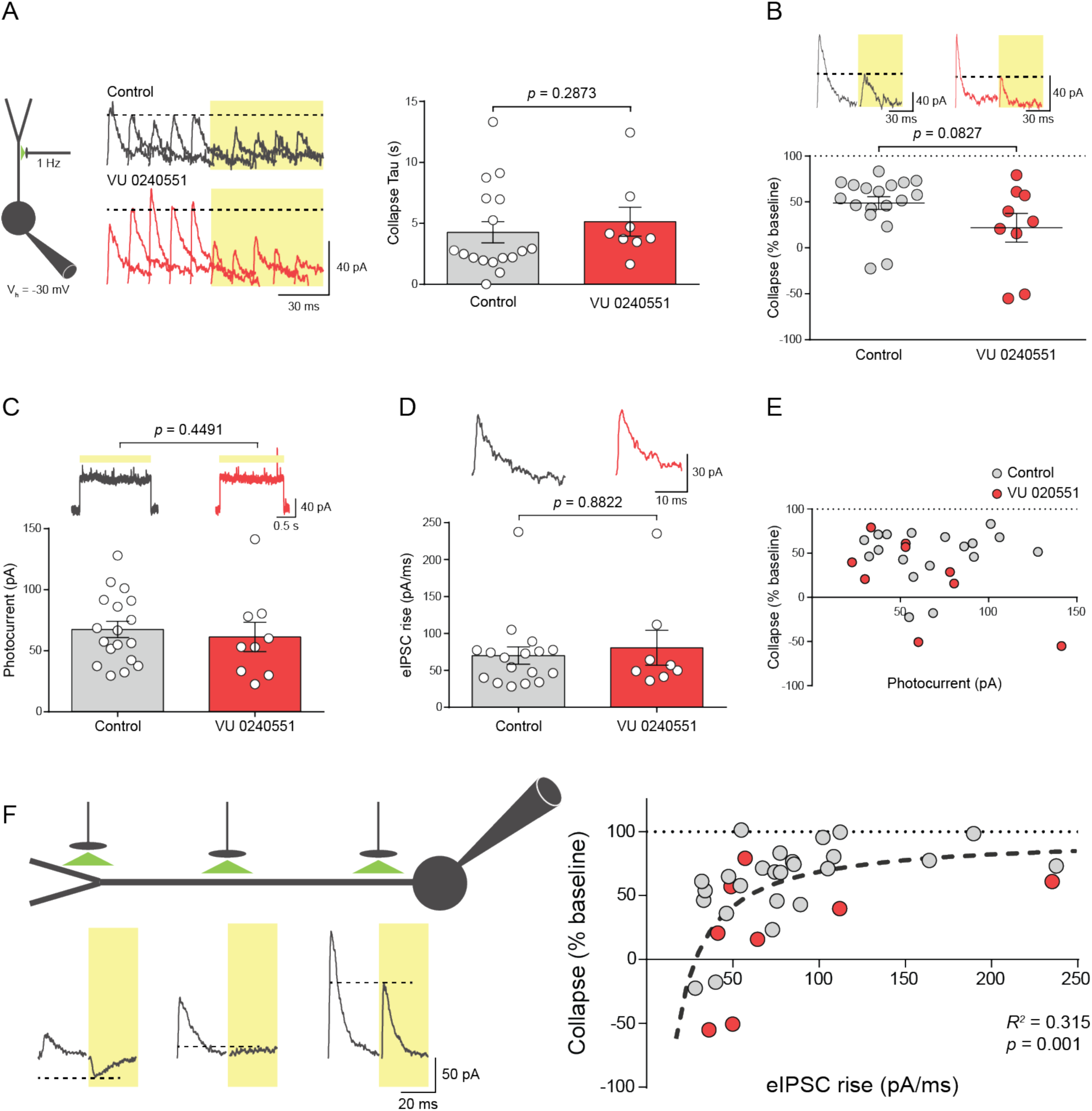
KCC2 does not set Cl^-^ buffering capacity in CRH^PVN^ neurons. **(A)** Left, schematic of stimulation and recording configuration. CRH^PVN^ neurons were voltage clamped at −30 mV and eIPSCs were elicited at 1 Hz. Middle, representative example of eIPSC currents before and during halorhodopsin activation in control cells and VU 0240551-treated cells. Right, VU 0240551 application does not alter the tau of current collapse (Mann-Whitney test, *U* = 49.0, *p* = 0.29). **(B)** VU 0240551 application does not alter the degree of eIPSC collapse caused by halorhodopsin activation (Mann-Whitney test, *U* = 47.0, *p* = 0.08). Inset: representative synaptic currents immediately before and immediately after light illumination in control cells and VU 0240551-treated cells. **(C)** The photocurrents generated in control and VU 0240551-treated cells were comparable (Mann-Whitney test, *U* = 66.0, *p* = 0.45). Inset: representative photocurrents in control cells and VU 0240551-treated cells. **(D)** The rates of eIPSC rise were indistinguishable between control and VU 0240551-treated cells (Mann-Whitney test, *U* = 65.0, *p* = 0.88). Inset: example eIPSCs elicited in control cells and VU 0240551-treated cells. Yellow bars represent 595 nm light illumination. **(E)** Relationship between photocurrent magnitude and degree of eIPSC collapse during halorhodopsin activation in control and VU 0240551-treated cells (no relationship, α = 0.05). **(F)** Left, synaptic currents immediately before and immediately after light illumination for eIPSCs with slow rise (left), moderate rise (middle), and fast rise (right). Currents displayed below cartoon neuron to indicate presumed synaptic location. Right, the relationship between the rate of eIPSC rise and the degree eIPSC collapse during Cl^-^ loading in control and VU 0240551-treated cells (inverse function, *R*^*2*^ = 0.32, *F*(1,31) = 14.2, *p* < 0.001).

Two observations support this idea: First, the photocurrent size was a poor predictor of the eIPSC collapse (*r* = −0.22, *p* = 0.20; **Figure 3e**), with 30 pA photocurrents generating similar Cl^-^ shifts as 130 pA photocurrents. Second, the location of synapses strongly influenced the collapse of Cl^-^ gradients during Cl^-^ loads (inverse fit, *R*^*2*^ = 0.32, *F*(1, 31) = 14.2, *p* < 0.001; **Figure 3f**). Synapses at distal locations (i.e. lower slope) were more susceptible to halorhodopsin-mediated Cl^-^ loading. In fact, at some distal synapses, halorhodopsin activation depolarized E_GABA_ past −30 mV, such that the eIPSC amplitudes switched in polarity (**Figure 3f**). Conversely, synapses at proximal locations were resistant to Cl^-^ rundown (**Figure 3f**), in many cases, failing to deviate from Cl^-^ setpoints despite high photocurrent amplitudes (data not shown). These data suggest that the susceptibility of GABA synapses on CRH^PVN^ neurons to Cl^-^ rundown, particularly during high Cl^-^ loads, is tightly regulated by synapse location, but not KCC2.

### KCC2 rapidly restores Cl^-^ setpoints in CRH^PVN^ neurons

KCC2 and other Cl^-^ homeostatic mechanisms may allow synapses to re-establish Cl^-^ gradients following deviations from setpoints. The recovery of Cl^-^ gradients after Cl^-^ loading is slowed by KCC2 inhibition^25,26,31^ and accelerated by KCC2 overexpression^43^. To determine the contributions of KCC2 in Cl^-^ setpoint restoration at CRH^PVN^ neuron inhibitory synapses, we measured the recovery of eIPSCs following halorhodopsin activation in the presence of the KCC2 inhibitor, VU 020551. In control cells, IPSCs recovered rapidly (**Figure 4a**) and exhibited properties consistent with KCC2-mediated Cl^-^ homeostasis. Specifically, the speed of eIPSC recovery was related to the size of the photocurrent, where a larger photocurrent initiated a faster homeostatic response (inverse fit, *R*^*2*^ = 0.57, *F*(1, 16) = 21.4, *p* = 0.0001; **Figure 4b**). This relationship resembles the disinhibition of KCC2 extrusion seen at higher Cl^-^ concentrations via with-no-lysine (WNK) protein kinase inhibition^44^ and the elevation of KCC2 cotransport thermodynamic driving force seen during larger setpoint deviations^4^. Blockade of KCC2 slowed the recovery of eIPSCs following photoactivation (control mean = 3.9 ± 0.89 s, *n* = 18, *N* = 11; VU 020551 mean = 15.8 ± 4.8 s, *n* = 9, *N* = 6; Mann-Whitney test, *U* = 28.0, *p* = 0.005; **Figure 4a**) and abolished the relationship between photocurrent and recovery speed (**Figure 4c**). Collectively, these observations suggest that KCC2 mediates rapid Cl^-^ setpoint restoration at CRH^PVN^ inhibitory synapses, and that this homeostasis is tuned to the strength of Cl^-^ loads.

**Figure 4.**
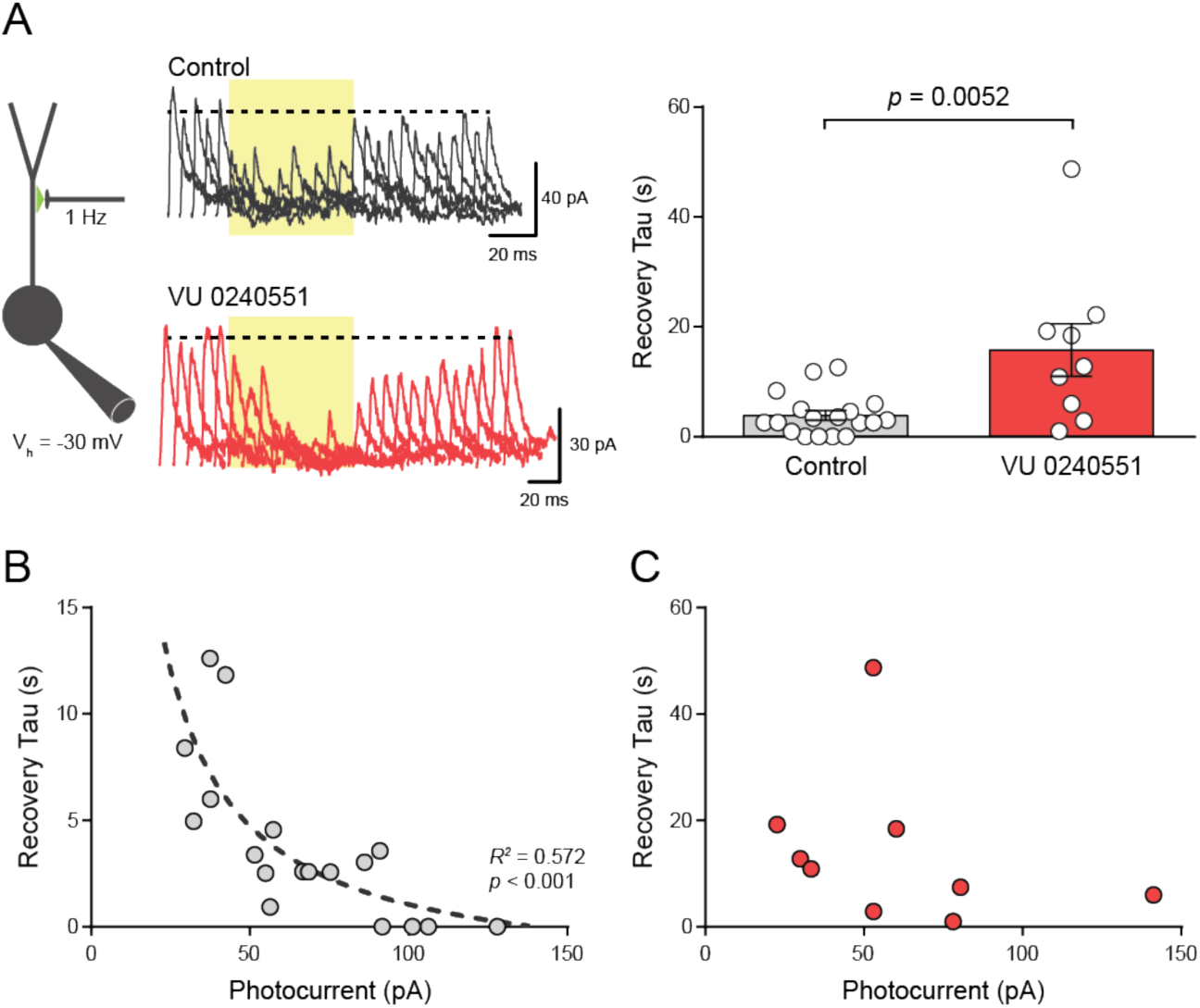
KCC2 rapidly restores chloride setpoints in CRH^PVN^ neurons. **(A)** Left, schematic of stimulation and recording configuration. CRH^PVN^ neurons were voltage clamped at −30 mV and eIPSCs were elicited at 1 Hz. Middle, representative eIPSC currents before, during, and after halorhodopsin activation in control and VU 0240551-treated cells. Right, VU 0240551 application slowed the recovery of eIPSCs following halorhodopsin activation (Mann-Whitney test, *U* = 28.0, *p* = 0.005). **(B)** The relationship between photocurrent magnitude and recovery of eIPSCs following Cl^-^ loading in control cells (inverse function, *R*^*2*^ = 0.57, *F*(1, 16) = 21.4, *p* < 0.0001). **(C)** The relationship between photocurrent and recovery of eIPSCs following halorhodopsin activation in cells treated with VU 0240551 (no relationship, α = 0.05). Yellow bars represent 595 nm light illumination.

### Acute stress slows Cl^-^ recovery in synaptic compartments

After characterizing synaptic Cl^-^ homeostasis in CRH^PVN^ neurons, and the role of KCC2 in these dynamics, we asked whether there were changes in Cl^-^ homeostasis across neuronal compartments or behavioural states. In the PVN, repetitive inhibitory synaptic input reduces spiking in CRH^PVN^ neurons from naïve animals but promotes spiking in CRH^PVN^ neurons from stressed animals^9^. A downregulation in KCC2 is thought to mediate this shift in GABAergic signalling. To directly test this idea, we stressed mice using a single 30-minute restraint stress and then measured Cl^-^ homeostasis at inhibitory synapses onto CRH^PVN^ neurons. The restraint stress protocol robustly and reliably elevated systemic CORT levels (naïve mean = 14.5 ± 2.6 ng/mL, *N* = 13; stress mean = 249.1 ± 15.2 ng/mL, *N* = 16; unpaired t-test, *t*(27) = 13.8, *p* < 0.0001). Immediately after the acute stress, brain slices were prepared and Cl^-^ homeostasis was probed. The recovery of synaptic eIPSCs following halorhodopsin photoactivation was slower in stressed animals compared to naïve animals (control mean = 3.9 ± 0.89 s, *n* = 18, *N* = 11; stress mean= 14.6 ± 4.1 s, *n* = 9, *N* = 7; Mann-Whitney test, *U* = 39.0, *p* = 0.03; **Figure 5a**). This indicates that a single bout of acute stress is sufficient to functionally impair Cl^-^ homeostasis.

**Figure 5.**
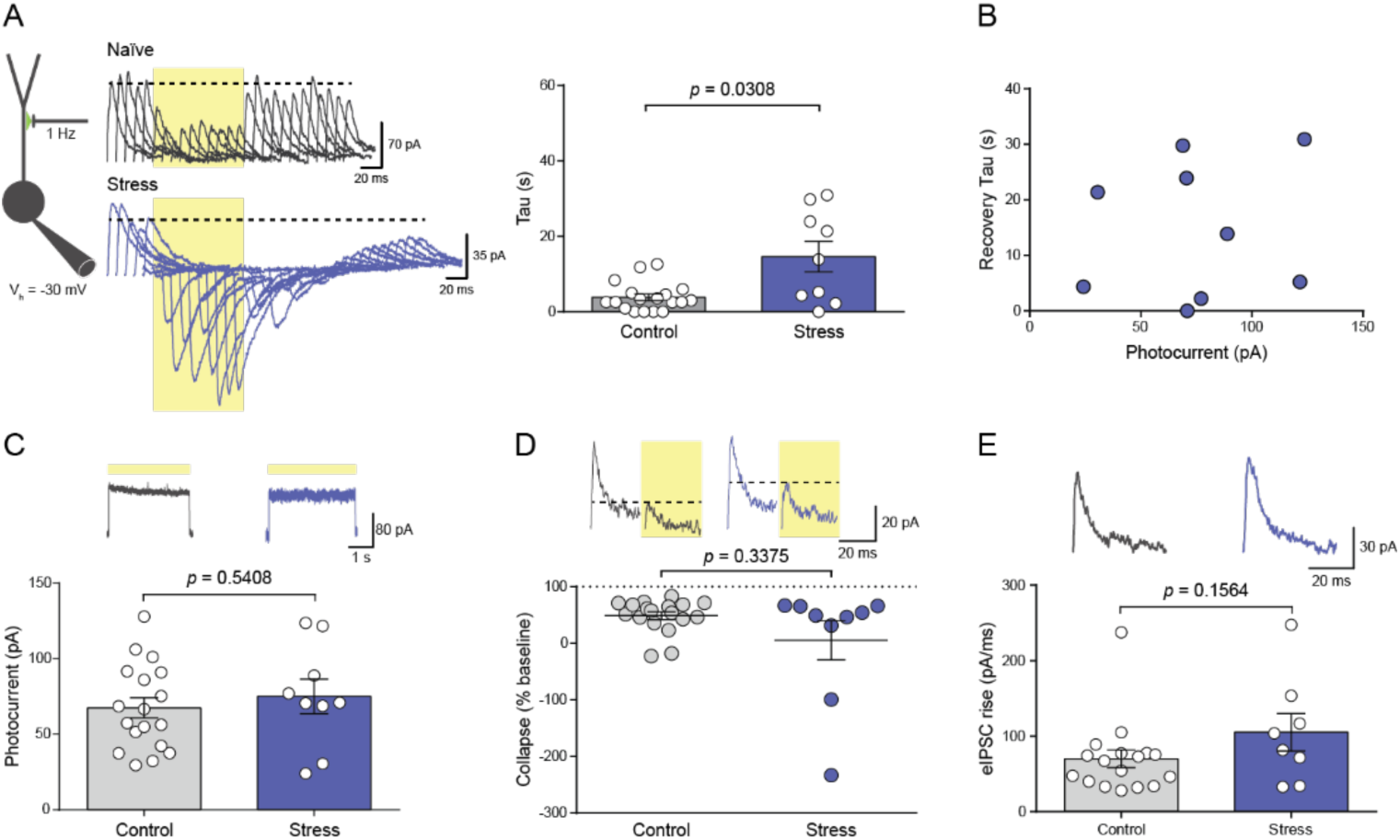
Stress slows synaptic chloride homeostasis in CRH^PVN^ neurons. **(A)** Left, schematic of stimulation and recording configuration. CRH^PVN^ neurons were voltage clamped at −30 mV and eIPSCs were elicited at 1 Hz. Middle, representative eIPSC currents before, during, and after halorhodopsin activation in cells from control and stressed mice. Right, the recovery of eIPSC amplitudes following halorhodopsin activation was slowed in cells from acutely stressed animals (Mann-Whitney test, *U* = 39.0, *p* = 0.03). **(B)** The relationship between photocurrent and recovery of eIPSCs following Cl^-^ loading in cells from stressed animals (no relationship, α = 0.05). **(C)** Photocurrents generated in cells from control and stressed animals were comparable (Unpaired t-test, t(25) = 0.62, p = 0.54). Inset: representative photocurrents in cells from control and stressed animals. **(D)** The degree of eIPSC shift during halorhodopsin activation is comparable in cells from control and stressed animals (Mann-Whitney test, *U* = 62.0, *p* = 0.34). Inset: representative synaptic currents immediately before and immediately after light illumination in cells from control and stressed animals. **(E)** The rates of eIPSC rise were indistinguishable between cells from control and stressed animals (Mann-Whitney test, *U* = 43.0, *p* = 0.16). Inset: example eIPSCs elicited in cells from control and stressed animals. Yellow bars represent 595 nm light illumination.

Again, the relationship between photocurrent and speed of recovery was abolished in cells from acutely stressed animals (**Figure 5b**). The effect of stress on Cl^-^ homeostasis was not due to differences in photocurrents (control mean = 67.5 ± 6.7 pA, *n* = 18, *N* = 11; stress mean = 75.1 ± 11.4 pA, *n* = 9, *N* = 7; unpaired t-test, *t*(25) = 0.62, *p* = 0.54; **Figure 5c**), Cl^-^ accumulation (control mean = 48.8 ± 6.9 %, *n* = 18, *N* = 11; stress mean = 5.3 ± 34.5 %, *n* = 9, *N* = 7; Mann-Whitney test, *U* = 62.0, *p* = 0.34; **Figure 5d**), or synaptic location (control mean = 70.2 ± 11.8 pA/ms, *n* = 18, *N* = 10; stress mean = 105.3 ± 24.9 pA/ms, *n* = 9, *N* = 7; Mann-Whitney test, *U* = 43.0, *p* = 0.16; **Figure 5e**). Taken together, KCC2 downregulation in CRH^PVN^ neurons during stress slows the restoration of Cl^-^ setpoints at inhibitory synapses.

### Cl^-^ homeostasis is maintained in somatic compartments during stress

Cl^-^ homeostasis in CRH^PVN^ neurons may also vary between cellular compartments. Inhibitory synapses onto PVN neurons innervate somatic and dendritic compartments, with dendritic synapses outweighing somatic synapses with a ratio of ∼3:1^45^. Somatic synapses often contain rapid Cl^-^ homeostatic mechanisms to maintain robust somatic inhibition, while dendritic synapses are usually more susceptible to Cl^-^ rundown^1,2^. To test if analogous segregations exist in CRH^PVN^ neurons or if differences can emerge during stress, we modified our optogenetic strategy to measure somatic Cl^-^ homeostasis in CRH^PVN^ neurons. Enhanced halorhodopsin was expressed in CRH^PVN^ neurons and then somatic GABA currents (I_GABA_) were tracked by voltage clamping cells at −30 mV and pressure applying GABA to the soma at 1 Hz. Similar to synaptic Cl^-^ homeostasis, somatic I_GABA_ amplitudes decayed to new steady state values during halorhodopsin activation, and then quickly recovered to baseline values after light termination (**Figure 6a**). The collapse and recovery of somatic I_GABA_ amplitudes were fit with mono-exponential functions to extract timing parameters; somatic I_GABA_ collapse occurred with a tau of 3.2 ± 0.22 s (monoexponential decay: *R*^*2*^ = 0.90 ± 0.02, **Figure 6b**) and somatic I_GABA_ recovery occurred with a tau of 5.0 ± 1.0 s (monoexponential association: *R*^*2*^ = 0.68 ± 0.10, **Figure 6c**). Somatic I_GABA_ amplitudes during halorhodopsin activation were 79.6 ± 2.5% of baseline values (**Figure 6b**). Using these parameters, we could characterize somatic Cl^-^ homeostasis in CRH^PVN^ neurons from naïve and acutely stressed animals.

**Figure 6.**
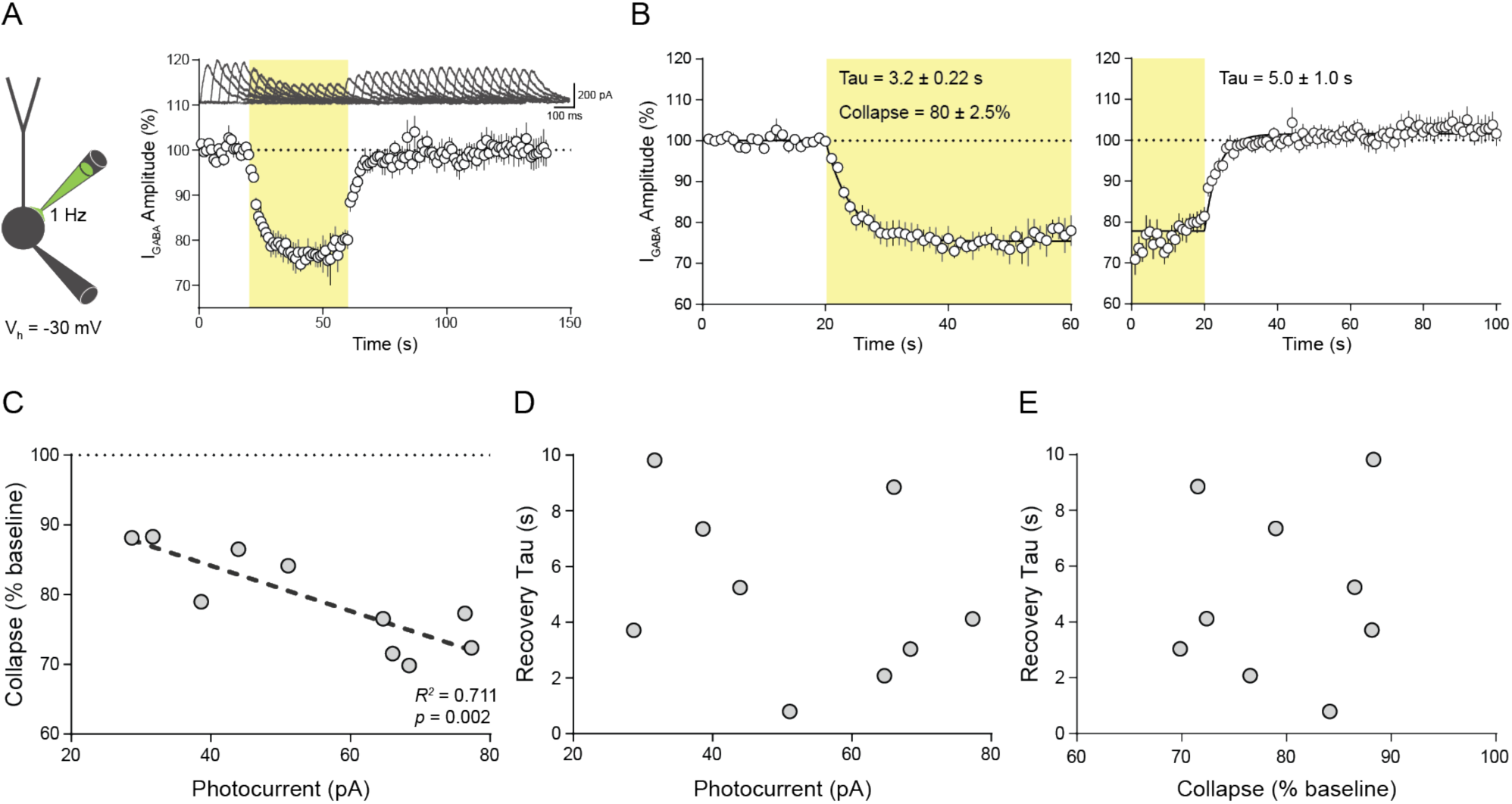
Somatic chloride homeostasis in CRH^PVN^ neurons. **(A)** Left, schematic of stimulation and recording configuration. CRH^PVN^ neurons were voltage clamped at −30 mV and I_GABA_ were elicited at 1 Hz. Right, average I_GABA_ amplitudes before, during, and after halorhodopsin activation, normalized to pre-light amplitudes. Inset, representative example of eIPSCs throughout experiment. **(B)** Left, average I_GABA_ amplitudes before and during light illumination, normalized to pre-light amplitudes. Data is fit with a monoexponential decay function (mean *R*^*2*^ = 0.90 ± 0.02) with an average tau of 3.2 ± 0.22 s. Right, average I_GABA_ amplitudes during and after light illumination, normalized to pre-light amplitudes. Data is fit with a monoeponential association function (*R*^*2*^ = 0.68 ± 0.10) with an average tau of 5.0 ± 1.0 s. **(C)** The relationship between photocurrent magnitude and degree of I_GABA_ collapse during halorhodopsin activation in control cells (linear function, *r* = −0.84, *R*^*2*^ = 0.71, *p* = 0.002). **(D)** The relationship between photocurrent magnitude and recovery of I_GABA_ following Cl-loading in control cells (no relationship, α = 0.05). **(E)** The relationship between degree of Cl^-^ collapse during light illumination and recovery of I_GABA_ following Cl-loading (no relationship, α = 0.05). Yellow bars represent 595 nm light illumination.

In naïve animals, somatic Cl^-^ homeostasis differed from synaptic Cl^-^ homeostasis in a few subtle ways. First, the average somatic I_GABA_ collapsed to 79.6 ± 2.5 % of baseline values (**Figure 6b**), with no currents reversing from outward to inward. Second, the collapse in somatic I_GABA_ was proportional to the size of the photocurrents (linear fit, *r* = −0.84, *R*^*2*^ = 0.71, *p* = 0.002; **Figure 6c**). Third, the recovery speed of somatic I_GABA_ following halorhodopsin activation was unrelated to the photocurrent magnitude (**Figure 6d**) or somatic I_GABA_ collapse (**Figure 6e**). These properties are consistent with the idea that somatic Cl^-^ gradients are resistant to Cl^-^ deviations^2^. Furthermore, the decoupling of recovery speed from photocurrent magnitude may indicate inherent differences between dendritic and somatic Cl^-^ homeostatic machinery.

The differences between somatic and synaptic Cl^-^ homeostasis segregated further after an acute stress. A single 30-minute restraint stress did not alter the recovery of somatic I_GABA_ following halorhodopsin activation (naïve mean = 4.9 ± 1.0 s, *n* = 9, *N* = 7; stressed mean = 12.8 ± 6.5 s, *n* = 10, *N* = 8; Mann-Whitney test, *U* = 35.0, *p* = 0.44; **Figure 7a**). This lack of effect was not explained by differences in I_GABA_ collapse during halorhodopsin activation (naïve mean = 79.4 ± 2.2 %, *n* = 9, *N* = 7; stressed mean = 79.4 ± 2.4 %, *n* = 10, *N* = 8; unpaired t-test, t(17) = 0.47, <u>p</u> = 0.65, **Figure 7b**) or photocurrents (naïve mean = 54.7 ± 5.8 pA, *n* = 9, *N* = 7; stressed mean = 51.8 ± 5.4 pA, *n* = 10, *N* = 8; unpaired t-test, *t*(20) = 0.37, *p* = 0.72, **Figure 7c**). One possible explanation for the discrepancy between synaptic and somatic homeostasis is that Cl^-^ diffusion from the patch pipette may limit our ability to detect somatic Cl^-^ dynamics. To show that somatic Cl^-^ homeostasis is sensitive to KCC2 manipulations, we lowered intracellular Cl^-^ to inhibit KCC2 function via WNK activation^44^ and to reduce the thermodynamic driving force of KCC2-mediated cotransport^4^. Indeed, lowering the Cl^-^ concentration of the internal solution from 12 mM to 4 mM decreased the recovery speed of somatic Cl^-^ currents following halorhodopsin activation (12 mM [Cl^-^]_i_ mean = 4.7 ± 0.97 s, *n* = 9, *N* = 7; 4 mM [Cl^-^]_i_ mean = 11.8 ± 2.5 s, *n* = 9, *N* = 9; unpaired t-test, *t*(16) = 2.5, *p* = 0.03; **Figure 7d**). Again, the recovery was not driven by changes in somatic I_GABA_ collapse (12 mM [Cl^-^]_i_ mean = 79.6 ± 2.5 %, *n* = 10, *N* = 7; 4 mM [Cl^-^]_i_ mean = 80.3± 3.4 %, *n* = 9, *N* = 9; Mann-Whitney test, *U* = 33.0, *p* = 0.53, **Figure 7e**) or photocurrents (12 mM [Cl^-^]_i_ mean = 54.7 ± 5.8 pA, *n* = 9, *N* = 7; 4 mM [Cl^-^]_i_ mean = 47.8 ± 10.3 pA, *n* = 9, *N* = 9; unpaired t-test, *t*(17) = 0.60, *p* = 0.56, **Figure 7f**). These results demonstrate that stress does not slow Cl^-^ homeostasis in the soma of CRH^PVN^ neurons and implies that dendritic and somatic inhibitory synapses are differentially regulated during stress.

**Figure 7.**
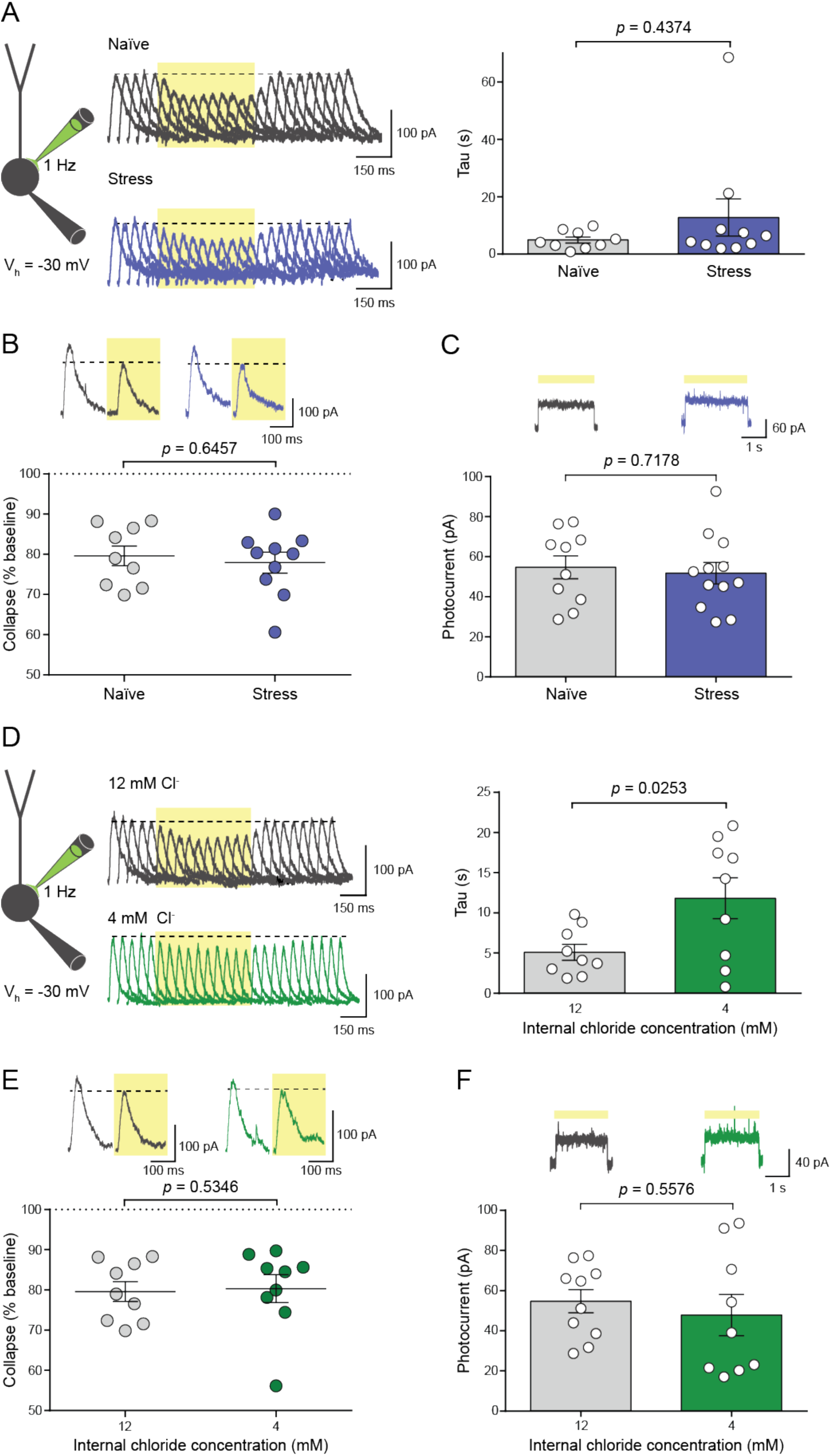
Stress does not alter somatic Cl^-^ homeostasis. **(A)** Left, schematic of stimulation and recording configuration. CRH^PVN^ neurons were voltage clamped at −30 mV and I_GABA_ were elicited at 1 Hz. Middle, representative I_GABA_ before, during, and after halorhodopsin activation in cells from control and stressed animals. Right, acute stress does not alter recovery of I_GABA_ amplitudes following Cl-loading (Mann-Whitney test, *U* = 35.0, *p* = 0.44). **(B)** The degree of I_GABA_ shift during halorhodopsin activation is comparable in cells from control and stressed animals (unpaired t-test, *t*(17) = 0.47, *p* = 0.65). Inset: representative I_GABA_ immediately before and immediately after light illumination in cells from control and stressed animals. **(C)** Photocurrents generated in cells from control and stressed animals were comparable (unpaired t-test, *t*(20) = 0.37, *p* = 0.72). Inset: representative photocurrents in cells from control and stressed animals. **(D)** Left, schematic of stimulation and recording configuration. CRHPVN neurons were voltage clamped at −30 mV and I_GABA_ were elicited at 1 Hz. Middle, representative I_GABA_ before, during, and after halorhodopsin activation in cells recorded with 12 mM [Cl-]_i_ and 4 mM [Cl-]_i_. Right, decreasing [Cl-]_i_ from 12 mM to 4 mM slows the recovery of I_GABA_ amplitudes following halorhodopsin activation (unpaired t-test, *t*(16) = 2.5, *p* = 0.03). **(E)** The degree of I_GABA_ shift during halorhodopsin activation is comparable in cells with 12 mM [Cl-]_i_ and 4 mM [Cl-]_i_ (Mann-Whitney test, *U* = 33.0, *p* = 0.53). Inset: representative I_GABA_ immediately before and immediately after light illumination in cells with 12 mM [Cl-]_i_ and 4 mM [Cl-]_i_. **(F)** Photocurrents generated in cells with 12 mM [Cl-]_i_ and 4 mM [Cl-]_i_ were comparable (unpaired t-test, *t*(20) = 0.60, *p* = 0.56). Inset: representative photocurrents in cells with 12 mM [Cl-]_i_ and 4 mM [Cl-]_i_.

## Discussion

Our results demonstrate that the cation-Cl co-transporter KCC2 rapidly restores Cl^-^ setpoints at GABA synapses onto CRH^PVN^ neurons. Furthermore, a decrease in KCC2 extrusion capacity following acute stress impairs Cl^-^ setpoint restoration at GABA synapses, but not at GABA_A_ receptors on the soma. These findings strongly support the idea that KCC2 functions primarily to maintain Cl^-^ setpoints and demonstrate that stress differentially modulates inhibitory synapses onto CRH^PVN^ neurons, such that dendritic synapses become more susceptible to Cl^-^ rundown and excitatory GABA signalling, while putative somatic synapses remain strongly inhibitory.

### Halorhodopsin and voltage clamp electrophysiology as a measure of Cl^-^ homeostasis

We used halorhodopsin and voltage-clamp electrophysiology to study how efficiently CRH^PVN^ neurons restore Cl^-^ driving force. Halorhodopsin was used to load Cl^-^ into cells and inhibitory synaptic currents or somatic GABA currents were used to track the resulting Cl^-^ dynamics. This approach has several advantages for the study of Cl^-^ regulation. First, by tracking the restoration of Cl^-^ setpoints, we directly measure Cl^-^ homeostasis and can make more conclusive statements about transporter function. Approaches that measure changes in E_GABA_ or E_IPSC_, in contrast, do not measure Cl^-^ homeostasis and can be misleading. For instance, blockade of KCC2 or NKCC1 can increase or decrease intracellular Cl^-^ concentrations depending on the initial concentration of Cl^-5^. In addition, Cl^-^ setpoints are highly sensitive to local impermeable anions and modifications to these anions can robustly shift Cl^-^ setpoints^4,5^. Our approach allows us to bypass these issues by directly measuring Cl^-^ dynamics. Second, halorhodopsin-mediated Cl^-^ loading does not interfere with voltage-clamp recordings of synaptic or extrasynaptic GABA_A_ currents. Alternative methods to load Cl^-^, such as repetitive synaptic stimulation or prolonged GABA application, can induce short-term plasticity^35^ or GABA_A_ receptor desensitization^2,34^, respectively. As short-term plasticity can differ between synaptic compartments^35^ and across behavioural states^46^, this interference can complicate data interpretation. Our approach avoids these issues by decoupling Cl^-^ measurements from Cl^-^ manipulations. Third, by tracking relative changes in Cl^-^ levels, we can use whole-cell patch clamp configurations instead of perforated patch clamp methods. This is a particular strength of our technique because gramicidin perforated patch clamp recordings are slow, low throughput, and difficult. In contrast, our recordings take less than three minutes to perform.

In our study, we measured Cl^-^ homeostasis across space and time using synaptic stimulation and GABA application. Future studies in the PVN or other brain regions can likely combine our optogenetic strategy with other optical methods to study more complex Cl^-^ heterogeneities. One possibility is to combine halorhodopsin-mediated Cl^-^ loading with channelrhodopsin-mediated synaptic stimulation to compare Cl^-^ homeostasis at synapses with genetically defined presynaptic cell types. A second possibility is to combine halorhodopsin with two-photon GABA uncaging to study the spatial organization of Cl^-^ homeostasis in cells with complex dendritic morphologies.

### Inhibition in the PVN during non-stress states

CRH^PVN^ neurons gate the endocrine and behavioural stress response. Activity in these cells drive the release of corticosteroids^19^ and the manifestation of stress-related behaviours^19–21^. To prevent these responses at inappropriate times, the PVN receives robust inhibitory inputs, compromising approximately 50% of all synaptic input^47^, that largely function to curtail CRH^PVN^ neuron activity^9,48^. During an acute stress, the synaptic input received by these neurons shifts to promote CRH^PVN^ activity^49–52^. A crucial part of this process is the relief of CRH^PVN^ neurons from robust inhibition. Stress circuitry accomplishes this task by downregulating KCC2 to promote Cl^-^ accumulation and excitatory GABA signalling^9^. Prior to our study, we did not fully understand how these changes shape Cl^-^ dynamics in CRH^PVN^ neurons.

Cation-Cl^-^ cotransport by KCC2 is thought to buffer Cl^-^ loads^8,9,23^ and restore equilibrium setpoints^2,8,9,23–26^. Our results demonstrate that KCC2 rapidly restores local Cl^-^ setpoints in CRH^PVN^ neurons, but do not influence the susceptibility of synapses to Cl^-^ rundown during large loads. Instead, synapse location, and presumably cell compartment volume^2^, play a larger role in Cl^-^ buffering and accumulation during loads. These results do not discount previous reports of enhanced Cl^-^ accumulation during KCC2 downregulation^8,9^. The restoration of Cl^-^ setpoints during repetitive synaptic stimulation is sufficient to explain these results. Our results instead place heavier emphasis on the role of neuronal geometry and synaptic properties on Cl^-^ dynamics during inhibitory signalling. Our results also reveal the relative speed of Cl^-^ homeostasis in CRH^PVN^ neurons. Although we measured Cl^-^ dynamics using whole-cell patch clamp configuration and cannot determine the precise kinetics of Cl^-^ homeostasis^2,32^, the restoration of dendritic and somatic Cl^-^ setpoints in CRH^PVN^ neurons was faster than analogous dynamics observed in hippocampal neurons^26^ and KCC2-overexpressing cortical neurons^43^ (also measured with whole-cell patch clamp). In many cases, Cl^-^ gradients recovered in sub-second timescales. We also found evidence that the speed of Cl^-^ restoration is tuned to the size of the Cl^-^ load. Taken together, CRH^PVN^ neurons contain efficient Cl^-^ homeostatic mechanisms that allow GABAergic input to maintain stable inhibitory signalling during naïve states.

### The functional organization of inhibition in the PVN changes during stress

We also provide direct evidence that stress slows KCC2-mediated Cl^-^ homeostasis at dendritic synapses onto CRH^PVN^ neurons, but not at somatic compartments. These results have several implications (**Figure 8**). First, slowing Cl^-^ homeostasis at dendritic synapses should cause major fluctuations in the strength and sign of inhibitory signalling. Using halorhodopsin, we showed that Cl^-^ loads can substantially shift the size and direction of inhibitory synaptic currents, with some distal synapses displaying E_IPSC_ depolarizations past −30 mV. During stress, large Cl^-^ deviations may go unchecked and thus create windows of time for excitatory GABA signalling^9^. Since GABA-mediated Cl^-^ influx is dependent on the driving force of Cl^-^, E_IPSC_ depolarization and excitatory GABA signalling may preferentially occur when CRH^PVN^ neurons are spiking, which may in turn facilitate further spiking or other depolarization-dependent events such as inhibitory or excitatory synaptic plasticity^53–55^. Second, the persistence of rapid somatic Cl^-^ homeostasis during stress suggests that inhibitory signalling onto somatic compartments remains stable during stress. If the framework proposed above is correct, then somatic inhibition may terminate CRH^PVN^ spiking to prevent runaway activity or to shape the timing of CRH^PVN^ activity. Indeed, fiber photometry recordings of CRH^PVN^ neurons suggest that many aspects of activity in these cells during stress are transient and time-locked to external events, such as movement-bouts^20,52^. Determining the roles of dendritic and somatic inhibition on CRH^PVN^ neuron physiology is an exciting future direction that will shape how we think about PVN network dynamics and stress.

**Figure 8.**
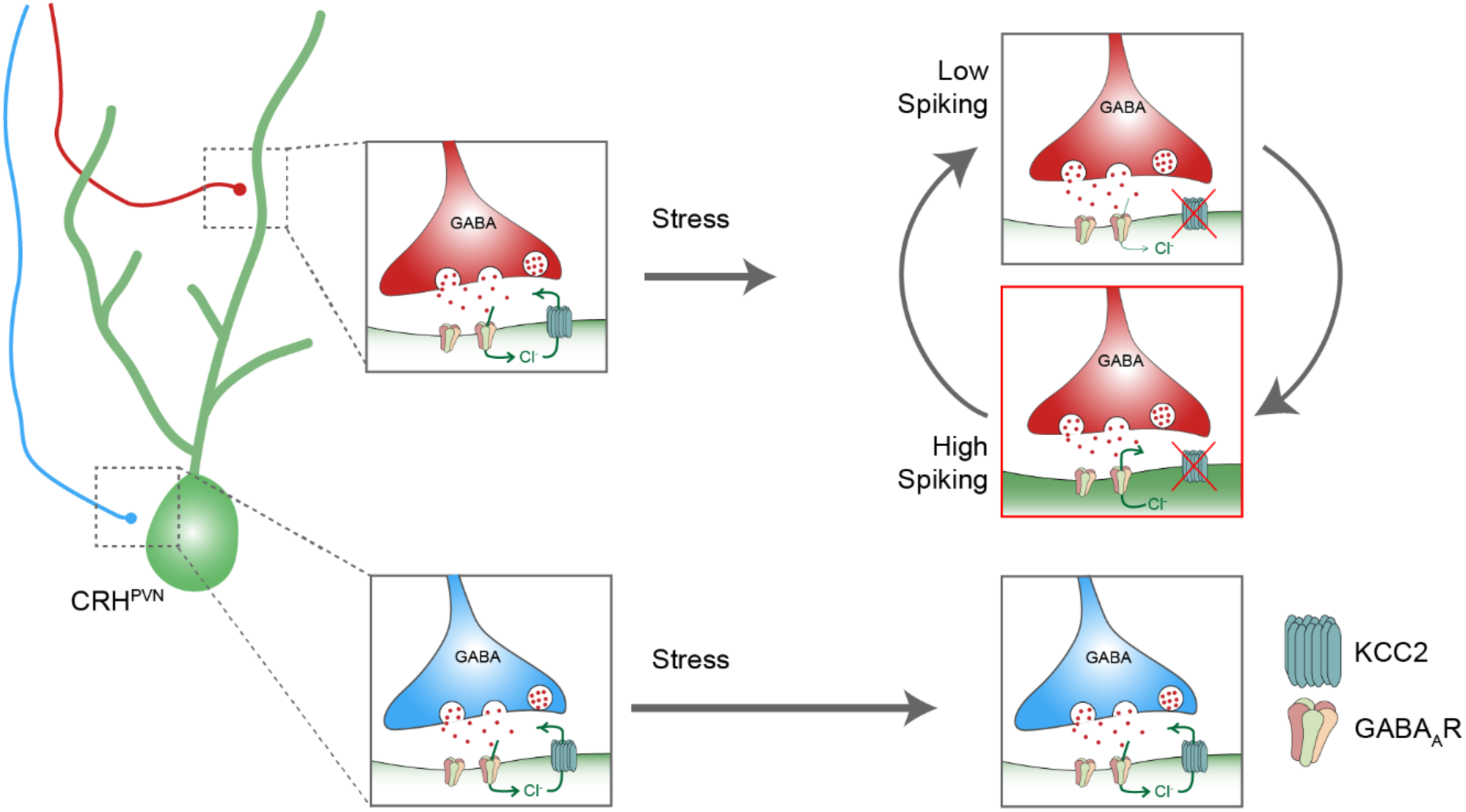
Dendritic and somatic GABA signalling onto CRH^PVN^ neurons. GABA signalling onto CRH^PVN^ neurons depends on the postsynaptic compartment (somatic vs dendritic) and the behavioural state (naïve vs stress). At somatic compartments (blue synapses), GABA signalling is strongly inhibitory during naïve and stressed states. This stability stems from the persistent action of KCC2 over time. In contrast, at dendritic compartments (red synapses), GABA signalling switches from inhibitory during naïve states to inhibitory and excitatory during stressed states. This bistability emerges because KCC2 is functionally downregulated during stress, causing a slowing in Cl^-^ setpoint restoration. Cl^-^ accumulation during high CRH^PVN^ spiking should promote excitatory GABA signalling, while Cl^-^ depletion during resting states should promote weak inhibitory GABA signalling.

What causes the slowing of Cl^-^ homeostasis in CRH^PVN^ neurons? Several pieces of evidence from this study and previous studies point towards a downregulation in KCC2. First, KCC2 blockade mimics the Cl^-^ dynamics seen during stress. Inhibiting KCC2 in CRH^PVN^ neurons from naïve animals depolarizes E_IPSC_ values^9^, slows Cl^-^ homeostasis, and abolishes the relationship between homeostasis and Cl^-^ load. Blocking KCC2 in cells from stressed animals has no effect on E_IPSC_ values^9^. Second, the Cl^-^ dynamics seen in these experiments were not affected by other homeostatic mechanisms. Blocking NKCC1 with furosemide does not shift E_IPSC_ values in CRH^PVN^ neurons from naïve animals^9^. Similarly, hyperpolarization-activated Cl^-^ channels are not activated by the voltage clamp values used during our experiments^56^. Despite this collection of evidence, KCC2 surface expression and dimerization are unaltered by acute stress^9^. A simple explanation for this discrepancy is that KCC2 proteins in CRH^PVN^ neurons are dynamically regulated by the WNK protein kinase during stress. WNK kinases activate a cascade of phosphorylation events that ultimately suppress the function of KCC2 without altering its expression^44,57^. The suppression of KCC2 by WNK kinases is regulated by Cl^-^ ions, such that elevations in Cl^-^ inhibit WNK kinases and disinhibit KCC2^44^. Work from our study indirectly supports the hypothesis that WNK-KCC2 interactions are heightened in CRH^PVN^ neurons during stress. In particular, the speed of homeostasis in cells from naïve animals was tightly coupled to intracellular Cl^-^ concentrations. Cl^-^ setpoint restoration was slowed when photocurrents were smaller or when the internal Cl^-^ concentration was lowered. This relationship mirrors the interactions between WNK and Cl^-^ concentration and suggests that KCC2 may be regulated by WNK-like pathways in CRH^PVN^ neurons. Conversely, during stress, the tight coupling between homeostasis and Cl^-^ is lost, suggesting the pathways involved in this regulation are somehow altered. As our evidence is correlational, further work is needed to unravel the molecular pathways controlling KCC2 function during stress.

## Conclusions

By using optogenetics and voltage clamp electrophysiology to directly measure Cl^-^ homeostasis, we have revealed complex spatiotemporal Cl^-^ dynamics in CRH^PVN^ neurons. These observations will allow us to develop more realistic predictions about the role of inhibition in CRH^PVN^ physiology and the interactions between stress and KCC2. This study emphasizes the power of using direct measurements of Cl^-^ homeostasis to study Cl^-^ regulation in the nervous system.

## Acknowledgments

We thank Ms. Cheryl Sank for technical assistance. This work was funded by a CIHR Foundation grant to JSB (FDN-148440). AL was supported by a CIHR Masters Scholarship. We are grateful to members of the Bains lab for many helpful discussions on earlier versions of this manuscript.

